# Droplet microfluidic screening to engineer angiotensin-converting enzyme 2 (ACE2) catalytic activity

**DOI:** 10.1101/2024.10.04.616690

**Authors:** Evelyn F Okal, Philip A. Romero, Pete Heinzelman

**Affiliations:** Department of Biochemistry, University of Wisconsin-Madison, Madison, WI, USA; Department of Biomedical Engineering, Duke University, Durham, NC, USA

**Keywords:** Amino acid racemase, Angiotensin-converting enzyme 2 (ACE2), Angiotensin-II, Directed evolution, High-throughput screening, Microfluidics, Peptidase, Protein engineering

## Abstract

**Background:** Angiotensin-Converting Enzyme 2 (ACE2) is a crucial peptidase in human peptide hormone signaling, catalyzing the conversion of Angiotensin-II to Angiotensin-(1-7), which activates the Mas receptor and elicits vasodilation, increased blood flow, reduced inflammation, and decreased pathological tissue remodeling. This study leverages protein engineering to enhance ACE2’s therapeutic potential for treating conditions such as respiratory viral infections, acute respiratory distress syndrome, and diabetes. Surrogate substrates used in traditional high-throughput screening methods for peptidases often fail to accurately mimic native substrates, leading to less effective enzyme variants. Here, we developed an ultra-high-throughput droplet microfluidic platform to screen peptidases on native peptide substrates. Our assay detects substrate cleavage via free amino acid release, providing a precise measurement of biologically relevant peptidase activity.

**Results:** Using this new platform, we screened a large library of ACE2 variants, identifying position 187 as a hotspot for enhancing enzyme activity. Further focused screening revealed the K187T variant, which exhibited a fourfold increase in catalytic efficiency (*k*_*cat*_*/K*_*M*_) over wild-type ACE2.

**Conclusions:** This work demonstrates the potential of droplet microfluidics for therapeutic peptidase engineering, offering a robust and accessible method to optimize enzyme properties for clinical applications.

## Background

Angiotensin-Converting Enzyme 2 (ACE2) is a peptidase that plays a central role in peptide hormone signaling pathways in humans. ACE2 cleaves the terminal phenylalanine residue from Angiotensin-II (Ang-II) to produce the Angiotensin-(1-7) peptide, which activates the Mas receptor (MasR). This activation results in downstream vasodilation, increased salutary blood flow, suppressed inflammation, and attenuated pathological tissue remodeling after acute injury [1,2]. These beneficial physiological responses have motivated researchers to pursue ACE2 as a therapeutic agent for treating respiratory viral infections [3], acute respiratory distress syndrome [4], and diabetes [5]. Our team, along with other researchers, is interested in using protein engineering techniques to enhance the activity and specificity of ACE2 to improve its therapeutic efficacy, specificity, and dosing [6].

Engineering peptidases such as ACE2 presents challenges related to high-throughput screening. Traditional methods for screening peptidase and protease activity often rely on chemically modified peptides with quenched fluorophores, which may not accurately mimic the shape and chemical properties of native substrates [7-9]. This discrepancy can lead to the identification of enzyme variants that preferentially hydrolyze the fluorogenic substrate over the native peptide substrate. Screening using native substrates is possible but typically requires lower throughput analytical techniques, such as liquid chromatography [6], which are time-consuming and labor-intensive. These screening challenges limit our ability to discover peptidase variants with enhanced activity on authentic peptide substrates.

In this study, we developed an ultra-high-throughput droplet microfluidic platform for screening peptidases on authentic peptide substrates. Our assay measures substrate cleavage by detecting the release of free amino acids, providing a more accurate readout of biologically relevant peptidase activity. Utilizing this advanced screening platform, we screened a large library of ACE2 variants for activity on the Angiotensin-II peptide substrate, generating extensive sequence-function data. Our findings identified position 187 as a critical hotspot for enhancing enzyme activity. Focused screening at this position led to the discovery of the K187T variant, which exhibited a fourfold increase in catalytic efficiency (*k*_*cat*_/*K*_*M*_) compared to wild-type ACE2. This work demonstrates the potential of droplet microfluidics to revolutionize the engineering of therapeutic peptidases, offering a robust and efficient approach to optimize ACE2 for clinical applications.

## Methods

### Fluorometric yeast-displayed ACE2 activity assay

We cloned the human ACE2 gene into a yeast surface display vector (Additional File 1 - **Fig S1**). Human ACE2- displaying yeast were cultured and induced as previously described [6] and 0.02 OD*mL of yeast cells were incubated in 150 μL of a 1:1 (vol/vol) mixture comprised of ACE2 reaction buffer (50 mM 2-(N-morpholino)ethanesulfonic acid (MES), 300 mM NaCl, 10 μM ZnCl2, 0.02% w/v Hen Egg Lysozyme (Sigma-Aldrich, St. Louis, MO) as carrier protein, pH 6.5) and L-amino acid quantitation kit (Sigma Aldrich, St. Louis, MO) reagent mix [10]. Reactions also contained Ang-II (Anaspec, Fremont, CA) at varying concentrations. Reactions were tumbled at 18 rpm on a tube rotator room temperature for ninety minutes, centrifuged to pellet yeast and supernatants withdrawn and transferred to black-walled, transparent bottom 96-well plates for fluorescence measurement using a Tecan Spark (Tecan, Morgan Hill, CA) platereader with excitation at 535 nm and emission at 587 nm.

### Microfluidic device fabrication

An initial layer of photoresist resin, SU-8 3010, was coated onto a mirrored silicon wafer (University Wafers, Boston, MA) and centrifuged at 1500 rpm to achieve 15 μm layer height. A photomask of the first layer of the microfluidic device was placed on the layer and 100 J/cm^2^ of UV light used to polymerize the features. The wafer was baked at 95 °C for 10 min to catalyze the polymerization. A second 25 μm layer of SU8-3025 was coated onto the wafer by spinning at 4000 rpm, and similarly polymerized with the second photomask to create the incubation line and baked again. Undeveloped photoresist was washed off with SU-8 developer (1-methoxy-2-propanol acetate, Fisher Scientific, Waltham, MA).

The wafer was then used to create a relief in un-polymerized polydimethylsiloxane (PDMS) (Dow Corning Sylgard® 184, 11:1 polymer:cross-linker ratio), which was then polymerized by baking at 75 °C. Inlet and outlet holes are punched with a 0.5 mm biopsy corer. The device was then thoroughly washed with isopropanol and double-deionized water and then plasma treated alongside a clean glass microscope slide, to which it was subsequently bonded. Prior to use, microfluidic channels were filled with Aquapel (Pittsburgh Glass Works, Pittsburgh, PA) to ensure hydrophobicity, and then baked for 10 min at 100 °C to vaporize any Aquapel left in the channels.

### Microfluidic analysis and sorting of ACE2 variants

A population of ACE2-displaying yeast were resuspended in the above MES reaction buffer, absent ZnCl_2_ to avoid precipitation, supplemented with 15% (v/v) Optiprep (Sigma), at a density of 10^4^ yeast per μL. This cell density was chosen to achieve one yeast cell per 10 drops to minimize double encapsulation events. Suspended yeast were loaded into 1 mL luer lock syringes, which were purged of air and fitted with luer-to-poly ethyl ethyl ketone (PEEK) tubing adapters. The cell syringe used PEEK tubing with 0.005” internal diameter, and all other syringes used 0.015” internal diameter PEEK tubing. Droplets containing yeast were generated at the co-flow drop maker junction. Both the yeast suspension and the assay reagents (25 μM Ang-II and amino acid quantitation kit enzyme reagents suspended in kit buffer at 2x concentration relative to the manufacturer’s protocol) were flowed into the device at 15 μL/h and were pinched into droplets by fluorinated oil (HFE 7500) containing 1% (wt/wt) PEG–perfluoropolyether amphiphilic block copolymer surfactant flowing at 100 μL/h.

The dropmaker was attached to a 30 cm PEEK incubation line that permitted the enzyme reactions to proceed for approximately 90 minutes before being reinjected onto the sorting chip. The droplets were sorted using electrocoalescence with an aqueous stream of Tris-EDTA buffer. A 532 nm laser was focused onto the channel just upstream of the sorting junction, each droplet was individually excited, and its fluorescence emission measured using a spectrally filtered PMT at 520 nm. A field-programmable gate array card controlled by custom LabVIEW code analyzed the droplet signal at 200 kHz, and if it detected sufficient fluorescence, a train of eight 180 V, 40 kHz pulses was applied by a high-voltage amplifier. This pulse destabilized the interface between the droplet and the adjacent aqueous stream, causing the droplet to merge with the stream via a thin-film instability, after which the droplet contents were injected into the collection stream via its surface tension. The sorted yeast cells were collected in a microcentrifuge tube and regrown in pH 4.5 Sabouraud Dextrose Casamino Acid media (SDCAA: Components per liter - 20 g dextrose, 6.7 grams yeast nitrogen base (VWR Scientific, Radnor, PA), 5 g Casamino Acids (VWR), 10.4 g sodium citrate, 7.4 g citric acid monohydrate) media [6] for subsequent analysis. Droplets were processed at 800-1000 Hz, and because the cell occupancy of the droplets was 10%, we thus analyzed 80-100 ACE2 variants per second.

### Mock sorting experiments to assess enrichment of active ACE2 variants

For the initial mock sorting experiments, we mixed ACE2-displaying yeast and negative control yeast displaying *L. lactis* EnlA, a protein that does not hydrolyze Ang-II, at a ratio of 1:10. This mixed population was loaded onto the microfluidic sorting system and the top 10% (by fluorescence signal) of the droplet population was isolated. The sorted yeast were cultured overnight in pH 4.5 SDCAA media at 30°C with shaking at 250 rpm for subsequent induction [6].

We quantified the activity of negative control, positive control, presorted, and post-sorted populations using a fluorogenic ACE2 substrate, Mca-Ala-Pro-Lys(Dnp) [Mca=(7-methoxycoumarin-4-yl)acetyl; Dnp=2,4-dinitrophenyl] (Enzo Life Sciences, Farmingdale, NY). Fluorogenic substrate reactions were carried out for ninety minutes in the above MES reaction buffer with addition of 15 μM substrate. Substrate hydrolysis was measured by analyzing reaction supernatants on a Tecan Spark platereader with excitation at 328 nm and emission at 393 nm.

### ACE2 mutational scanning

For ACE2 random mutant library construction, the GeneMorph II Kit (Agilent Technologies, Santa Clara, CA) was employed using the wild type ACE2 display plasmid as template and respective forward and reverse primers CDspLt (5’-GTCTTGTTGGCTATCTTCGCTG-3’) and CDspRt (5’-GTCGTTGACAAAGAGTACG-3’). Error prone PCR products from the GeneMorph random mutagenesis reactions were digested NheI to MluI and ligated into the VLRB.2D-aga2 vector [6] digested with these same enzymes and transformed into yeast display *Saccharomyces cerevisiae* strain EBY100 as described [11]. The transformed library was grown in at 30°C and 250 rpm for two days. An aliquot of the transformant pool was plated on SD -Trp agar to quantify the number of library transformants. Plate counts showed that the yeast-transformed library contained ∼2*10^5^ unique clones; sequencing of fifteen randomly chosen clones from the library showed an average coding mutation rate of 1.8 per gene.

For induction of ACE2 mutant library display a 5 mL Sabouraud Galactose Casamino Acid (SGCAA: Components per liter - 8.6 g NaH_2_PO_4_*H_2_O, 5.4 g Na_2_HPO_4_, 20 g galactose, 6.7 g yeast nitrogen base, 5 g Casamino Acids) was started at an optical density, as measured at 600 nm, of 0.5 and shaken overnight at 250 rpm and 20°C. After overnight induction approximately 2*10^6^ yeast cells were harvested by centrifugation, washed once in pH 7.4 Phosphate Buffered Saline (PBS) containing 0.2% (w/v) bovine serum albumin (BSA) and encapsulated in amino acid quantitation kit reagent-containing droplets as described above. Three independent droplet sorts were performed: one sort of 10^6^ yeast with isolation of the top 5% of the yeast population, one sort of 7*10^5^ yeast with isolation of the top 20%, and a third sort of 10^6^ yeast with isolation of the top 20%; all sorts featured 25 μM Ang-II as ACE2 hydrolysis substrate.

Yeast cells isolated during sorting were cultured overnight in 5 mL of low pH SDCAA media at 30°C with shaking at 250 rpm and the following morning 1/10th of the culture volume was expanded in low pH SDCAA to an OD of 0.1, shaken at 30°C and 250 rpm, and harvested by centrifugation after the OD had reached 0.4. In parallel, yeast culturing, at 30 mL scale, and harvesting was carried out for the pre-sort random mutant library yeast populations used as inputs for the three respective sorts. ACE2 mutant yeast display plasmids were rescued using the ZymoPrep Yeast Plasmid Miniprep II kit (Zymo Research). Rescued plasmids were amplified via electroporation into 10G Supreme *E. coli* (Lucigen, Madison, WI) with subsequent culturing and DNA harvest as previously described [11].

For NGS analysis, amplified plasmid DNA from the three respective pre- and post-sort yeast populations were digested in separate reactions using PstI and XhoI restriction enzymes. Digested plasmids were run in a 1.2% agarose gel and the ACE2 band was excised and purified using the Zymo Gel Extraction kit (Zymo Research, Orange, CA). Purified DNA was submitted to the University of Wisconsin-Madison Biotechnology Center DNA Sequencing Facility and a sequencing library was prepared using the NEBNext Ultra II kit (New England Biolabs, Beverly, MA). Samples were sequenced using an Illumina NovaSeq6000 (Illumina, San Diego, CA) with 2×150 bp reads.

The reads from the Illumina FASTQ files were mapped to the wild type human ACE2 gene using Bowtie2 [9] and translated to amino acid sequences. Mutations observed fewer than ten times were discarded prior to continuing analysis. We used the presort and postsort mutation frequency data to calculate log enrichment values for each observed amino acid substitution. While we observed large variability in mutational enrichment across the three replicates, we focused on mutations with large enrichment values across the three replicates to identify the six ACE2 mutants: T20Ile, K187Q, R219K, W328R, Q340R and T371M were chosen for recloning into the yeast display plasmid and assessment of Ang-II hydrolyzing activity.

### Evaluation and directed evolution of lead ACE2 mutants identified by NGS data analysis

The above six ACE2 mutants were recloned into the yeast display plasmid and Ang-II hydrolysis assays, which featured 25 μM Ang-II as substrate and used liquid chromatography (LC) to quantify yeast-displayed ACE2 activity, were carried out as described [6] with the exception that assays were conducted with a range of numbers of yeast (0.02, 0.006 and 0.002 OD*mL) to ensure that activity differences across yeast variants would be fully broken out. The ACE2 position 187 site-directed mutant library was constructed by using overlap extension PCR to introduce a degenerate NNB codon and the resultant mutant library was DNA amplified and transformed into yeast display strain EBY100 as previously done for site-directed ACE2 mutant libraries containing substitutions at other positions [6]. Liquid cultures for fifty colonies from the position 187 site-directed mutant library transformant synthetic dropout -Trp agar plates (Teknova, Hollister, CA) were grown in SDCAA media and induced in SGCAA media [6]. Yeast-displayed ACE2 mutant activity was quantified by incubating yeast with 25 μM Ang-II as described [6] and analyzing reaction products by LC.

For flow cytometric quantification of wild type and position 187 mutant ACE2 yeast surface display levels, yeast colonies were picked into 4 mL of low pH SDCAA and grown overnight at 30°C with shaking at 250 rpm. Cultures were induced in 5 mL of SGCAA overnight at 20°C with shaking at 250 rpm; induction starting OD was 0.5. After induction yeast were harvested by centrifugation and washed in PBS/0.2% BSA and 3*10^5^ yeast were tumbled on a tube rotator at 18 rpm for two hours at 4°C in 300 μL of PBS/0.2% BSA containing 3 μg/mL anti-*myc* IgY (Aves Labs, Davis, CA). Following this primary labeling, yeast were washed in PBS/0.2% BSA and rotated at 18 rpm in 300 μL of same buffer containing 2 μg/mL Alexa488-conjugated goat anti-chicken IgG (Jackson ImmunoResearch, West Grove, PA) for one hour at 4°C. Yeast cells were washed in PBS/0.2% BSA and resuspended in ice cold PBS for analysis on a Fortessa (Becton Dickinson) flow cytometer in the UW-Madison Biochemical Sciences Building.

### Soluble production and Ang-II hydrolysis assay of ACE2 position 187 mutants

Plasmid pcDNA3-sACE2(WT)-8his [6], which encodes human ACE2 residues 1-615 with native human codon representation and puts secretion of the gene product, with a C-terminal His8 tag, from mammalian cells under control of the cytomegalovirus CMV promoter, was used as template for creation of ACE2 genes carrying mutations at position 187. Mutant genes (K187T and K187Q) were constructed by overlap extension PCR, ligated into pcDNA3-sACE2(WT)-8his, and transformants sequenced as described [6].

Transfections of plasmid DNA for wild type ACE2, the two K187 mutants, and a previously studied enhanced T371L/Y510Ile ACE2 double mutant [6] into Human Embryonic Kidney 293T cells (Product #CRL-3216, ATCC, Manassas, VA) were performed in duplicate and Ni-NTA purification of the secreted ACE2 proteins from cell culture media supernatant carried out as previously done for wild type and a panel of ACE2 mutants [6].

Purified proteins were buffer exchanged into 25mM Tris-HCl, pH 7.5 containing 10 μM ZnCl_2_ using Zeba Spin desalting columns (Fisher Scientific) and SDS-PAGE analysis of purified ACE2s was carried out using a LifeTech Mini Gel Tank electrophoresis system. Approximately 1.5 μg of purified, desalted ACE2 was loaded into each well of a Novex 4-20% Tris-Glycine gel and electrophoresis was performed at 200 volts for 45 minutes. 0.75 μg, 1.5 μg and 0.45 μg of BSA were loaded into separate wells of the gel to aid in determining ACE2 protein concentration by densitometry. PageRuler Unstained Protein Ladder (Fisher Scientific) was used as a molecular weight standard. Gels were stained with Simply Blue SafeStain (Invitrogen) following electrophoresis, photographed, and densitometry of gel photos was performed with GelQuant.NET software (Biochem Lab Solutions, San Francisco, CA) to determine ACE2 sample protein concentrations.

Ang-II substrate hydrolysis assays were carried out at room temperature and 150 μL scale in 96-well plates using 50 mM MES, 300 mM NaCl, 10 μM ZnCl_2_, 0.01% (vol/vol) Brij-35 (Sigma-Aldrich) as reaction buffer. Reactions were halted by addition of 15 μL of 1M EDTA solution, pH 8, after intervals, which ranged from eight minutes to thirty minutes, during which less than 15% of the input peptide substrate had been hydrolyzed. ACE2 was added to all reactions at 700 pM. Moles of product formed in hydrolysis reactions used for ACE2 activity and specificity profiling were determined by comparing area under the curve for LC chromatograms corresponding to these reactions to areas under the curve observed for control reactions in which > 95% input substrate conversion was achieved by incubation of substrate with 25 nM purified wild type ACE2 for seventy-five minutes. Elution times for peptide substrate inputs and hydrolysis products were determined by comparing chromatograms for the 25 nM ACE2 commercial standard reactions to input peptide reaction mixtures to which no ACE2 was added. LC analysis injection volumes and mobile phase gradient parameters were as previously described [6] and kinetic parameters for Ang-II hydrolysis were determined by fitting initial rates data using K_m_V_max_ Tool Kit online software (http://www.ic50.tk/kmvmax.html).

## Results

### A microfluidic platform for screening peptidases on native peptide substrates

Droplet microfluidics allows for the compartmentalization of biological assays in picoliter-scale water-in-oil emulsions, enabling ultra-high-throughput analysis of cells and biomolecules [9]. This method has significant potential for enzyme engineering, providing a platform to evaluate massive libraries of enzyme variants quickly and efficiently while using minimal reagents. We developed a microfluidic screening platform to screen ACE2 and other peptidases on peptide substrates (**Fig 1a**). Our workflow starts with a library of ACE2 variants displayed on the surface of yeast; individual yeast cells are encapsulated in microdroplets with the Angiotensin-II peptide and reagents to detect substrate hydrolysis. We then incubate the droplets for 60-90 minutes in an integrated delay line, allowing the ACE2 enzymes to cleave the substrate and for assay fluorescence to develop. The droplets are then reinjected onto a sorting chip where individual droplet fluorescence is measured, droplets showing high ACE2 activity are sorted, and the corresponding genes are collected and analyzed using Illumina sequencing.

**Figure 1:**
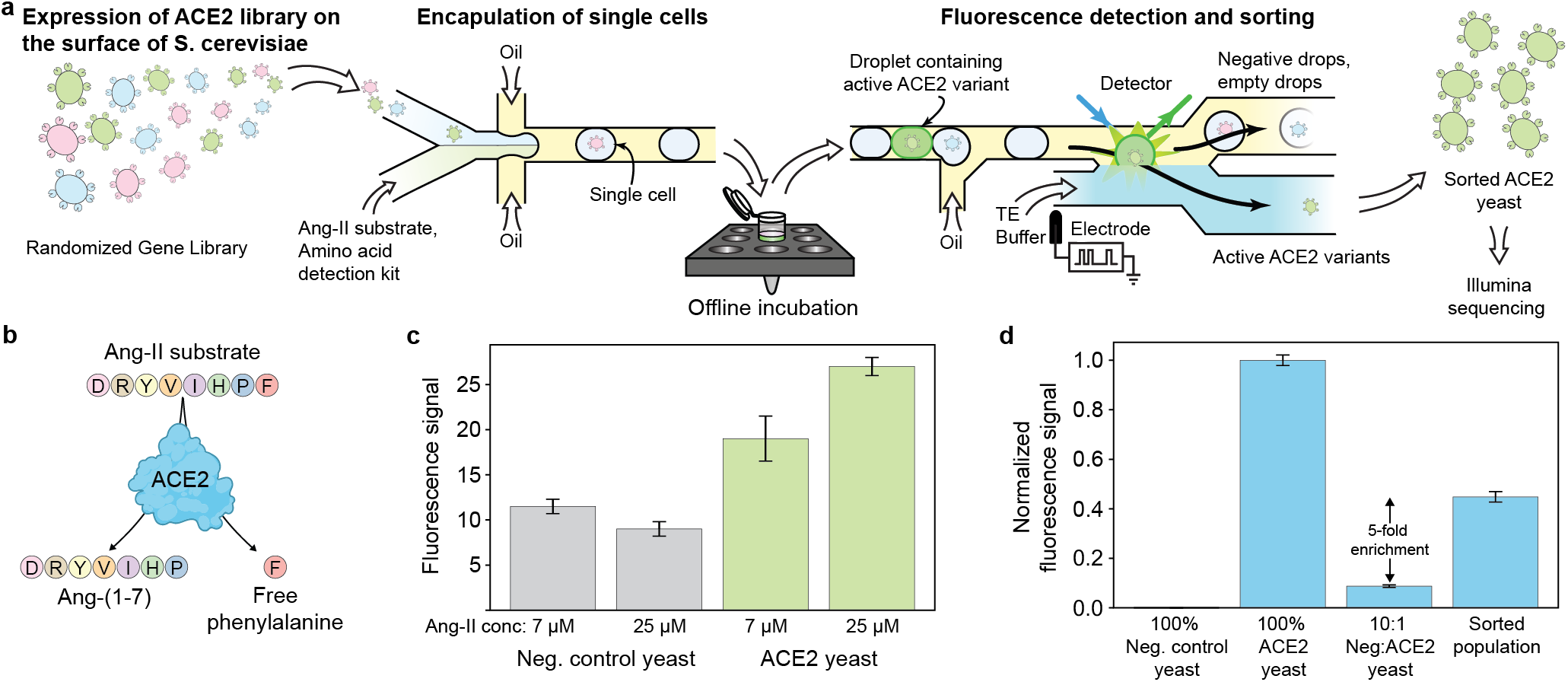
An ultra-high-throughput ACE2 screening platform. (**a**) An overview of our microfluidic screening system. Yeast displaying ACE2 variants are encapsulated in droplets with the Ang-II substrate and a phenylalanine detection kit. The droplets are incubated offline, reinjected onto a sorting chip, analyzed for fluorescence signal, and active ACE2 variants are sorted and collected for downstream analysis. (**b**) Ang-II cleavage by ACE2 releases free phenylalanine, which is detected using a commercial amino acid detection kit [10]. (**c**) Our ACE2 assay can reliably detect Ang-II cleavage by ACE2-displaying yeast. Error bars represent one standard deviation for triplicate measurements. (**d**) A mock sorting experiment where negative and positive control ACE2 yeast are combined at a 10:1 ratio, put through the full microfluidic screening pipline, and the pre/post-sort populations are analyzed for ACE2 activity. The microfluidic system can enrich active ACE2 fivefold. Error bars represent one standard deviation for triplicate measurements.

To address the challenge of screening peptidase activity on native substrates, we developed a coupled enzyme assay that detects the free phenylalanine released from Ang-II cleavage. We used a commercial kit [10] that detects amino acids using an ensemble of enzymes to reduce resazurin to resorufin and produce a fluorescence signal. We tested this assay in bulk using yeast-displayed ACE2 and negative controls (**Fig 1c**). Reaction time is a particularly important consideration due to the time dependent increase in background fluorescence signal that occurs due to spontaneous reduction of resazurin to the fluorescent resorufin compound and the assay signal saturation phenomenon that occurs due to the finite amounts of both Ang-II and resazurin present in the reaction mixture. The highest observed assay signal-to-background ratio was achieved for ninety-minute reactions containing 25 μM Ang-II. The assay can reliably detect ACE2 activity, ensuring that the detected activity accurately reflects ACE2’s ability to process native substrates, thereby providing reliable data for further engineering efforts.

We next evaluated our microfluidic system’s ability to enrich active ACE2 variants from a mixed population. We performed a mock sorting experiment by combining active ACE2 yeast with a tenfold excess of negative control yeast (**Fig 1d**). We ran this mixed control population through our microfluidic system and sorted droplets with high fluorescence values. We then measured the total ACE2 activity of the initial population and the sorted population, comparing them to pure ACE2 and negative control populations to estimate the proportion of active ACE2 yeast in each population. The initial population had 9% ACE2 activity, consistent with our 1:10 loading, while the sorted population had 45% ACE2 activity, indicating a fivefold enrichment of active ACE2. These results demonstrate our platform’s capability to screen large libraries of ACE2 variants and identify those with enhanced activity on native Ang-II substrates.

#### Mutational scanning of ACE2 to discover activity-boosting mutations

We applied our high-throughput microfluidic platform to systematically scan for mutations that could enhance ACE2 activity on Ang-II. Using error-prone PCR, we introduced random mutations throughout the ACE2 peptidase domain gene, ensuring broad coverage of potential activity-boosting mutations. These libraries contained 1.3 amino acid substitutions per variant. We used the microfluidic system to screen the ACE2 library in triplicate. For each screening run, we analyzed 650k - 1M variants and sorted 130k - 200k (top ∼5-20%) active ACE2 variants. The sorted yeast cells were regrown, and their ACE2 genes were extracted for downstream DNA sequencing analysis. We then sequenced the initial (presorted) library and the sorted library for each of the three replicates.

Using the sequence counts from the Illumina data, we calculated the log enrichment for individual amino acid substitutions. We found considerable variability between the replicates, indicating potential issues with the droplet assay or the microfluidic sorting (see Discussion). To robustly identify activity-boosting mutations, we focused on mutations that showed beneficial effects across all three replicates. We tested candidate mutations T20I, K187Q, R219K, W328R, Q340R, and T371M using a clonal yeast display assay [6] with LC readout of Ang-II cleavage (**Fig 2**). Most of the variants had decreased activity, but K187Q showed 1.5x greater activity than wild-type ACE2.

**Figure 2:**
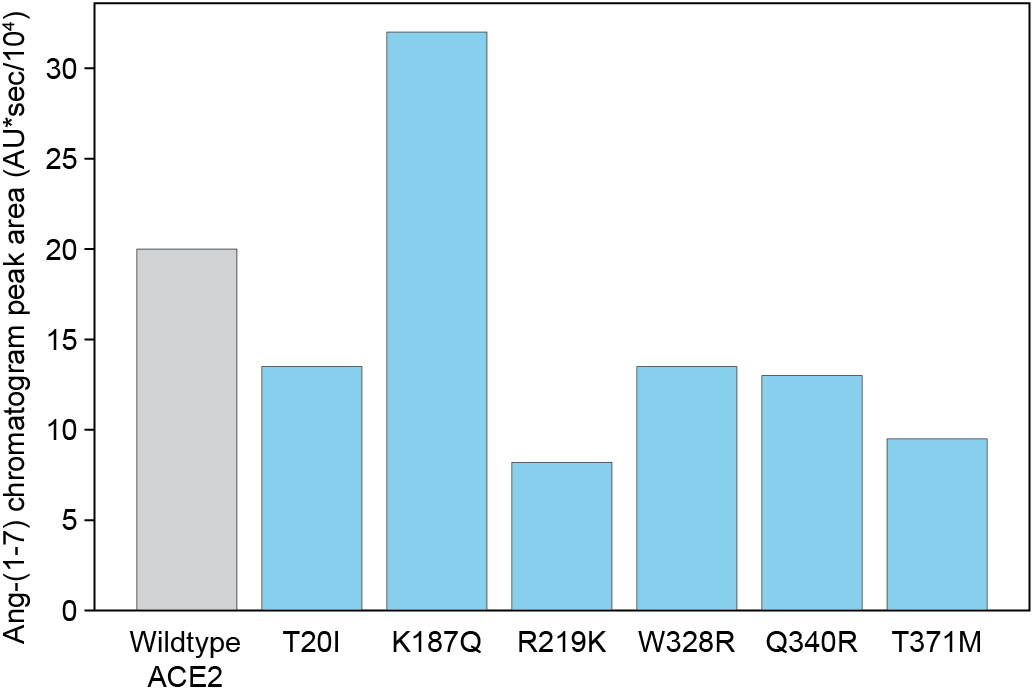
ACE2 variants prioritized by high-throughput mutational scan. Variants were displayed on yeast and assayed using the Ang-II substrate with liquid chromatography readout.

### Lysine 187 as a mutational hotspot to enhance ACE2 catalytic activity

We identified the K187Q mutation from our mutational scan and followed up on this result by using saturation mutagenesis at position 187 to test all possible amino acid substitutions. We generated an NNB library at position 187 and screened 50 variants using a clonal yeast display assay with LCMS readout of Ang-II cleavage. This screening revealed multiple amino acid substitutions that enhanced total ACE2 activity, including K187T, which exhibited over 2x higher total activity than wild-type ACE2 (**Fig 3a**). The observed increases in total activity could be attributed to either increased protein expression/display or intrinsic catalytic activity of the enzyme. To determine the cause, we performed flow cytometry studies to measure the protein display levels of the K187 mutants. We found that all variants displayed at levels similar to each other and wild-type ACE2 (Additional File 1 - **Fig S2**), suggesting that the observed differences in total activity are due to improvements in enzyme *k*_*cat*_ and/or *K*_*M*_ rather than variations in enzyme copy number on the yeast surface.

**Figure 3:**
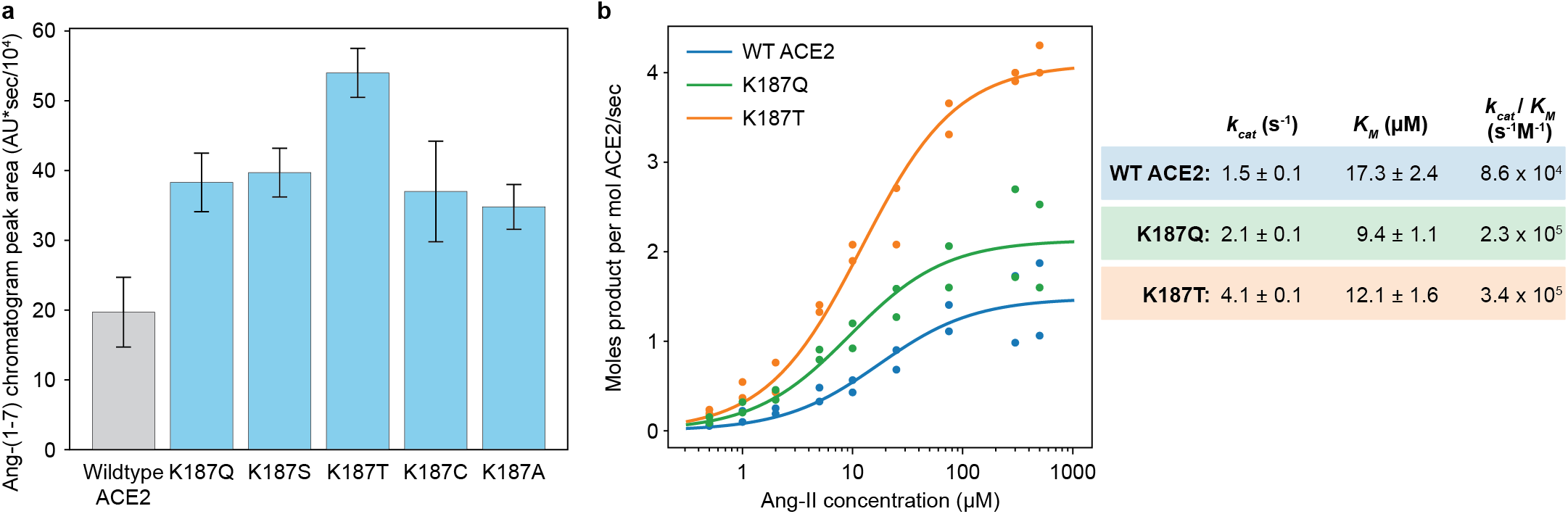
Mutations at position 187 enhance ACE2 catalytic activity. (**a**) Catalytic activity of K187 mutants. Multiple amino acid substitutions at position 187 enhance ACE2 activity. Error bars represent one standard deviation for triplicate measurements. (**b**) Initial rate plots of ACE2 and K187Q/T mutants. The K187 mutants have increased *k*_*cat*_ and decreased *K*_*M*_ relative to wildtype (WT) ACE2.

We performed Michaelis-Menten kinetic measurements to further explore the catalytic properties of these enzymes. We cloned the K187Q and K187T mutants into an ACE2 mammalian cell secretion vector, transiently transfected human embryonic kidney (HEK) cells, and purified the secreted ACE2 proteins from culture supernatant [6]. We performed initial reaction rate measurements across a broad range of Ang-II concentrations and found the K187Q and K187T mutants possess greater activity than wild type ACE2 for a broad range of Ang-II concentrations (**Fig 3b**). We fit the Michaelis-Menten parameters and found both K187Q and K187T have improved k_cat_ and K_m_. K187T had a 3.6-fold increase in catalytic efficiency (*k*_*cat*_/*K*_*M*_) compared to wild-type ACE2. The post-purification expression yields for the K187 mutants were either identical or similar to those obtained for wild type ACE2 (Additional File 1 - **Fig S3**).

## Discussion

We created an ultra-high-throughput droplet microfluidic system designed to screen peptidases using authentic peptide substrates. Our assay detects substrate cleavage through the release of free amino acids, yielding a precise measurement of biologically relevant peptidase activity. Using this screening platform, we analyzed a large library of ACE2 variants for their activity on the Angiotensin-II peptide substrate, which allowed us to discover activity boosting mutations. Our results pinpointed position 187 as a key site for increasing ACE2 catalytic activity. Saturation mutagenesis at this location uncovered the K187T variant, which demonstrated a fourfold increase in catalytic efficiency (*k*_*cat*_/*K*_*M*_) over wild-type ACE2. This study showcases the potential of droplet microfluidics to enable peptidase engineering for broad applications in industry and medicine.

Our microfluidic peptidase screening system is highly generalizable, uses commercially available materials, can be implemented with only basic microfluidic screening capabilities, and can be broadly applied to other medically relevant peptidases. Our approach relies on yeast surface display, which can reliably express large mammalian proteins with posttranslational modifications such as disulfide bonds and glycosylation. Peptidase enzymes such as microbial aminopeptidases used in food and pharmaceutical processing [12] and carboxypeptidases that play important roles in blood clotting [13] all cleave peptides to produce free amino acids, which can be detected with a commercial amino acid detection kit [10]. The results obtained also provide a foundation for extrapolating our method to the study and engineering of amino acid racemases [14] and enzymes used for synthesis of noncanonical amino acids [15]. The combination of a robust protein expression system, generalizable amino acid detection methods, and ultra-high-throughput droplet microfluidic screening that we have brought together in this work will open new avenues in therapeutic enzyme engineering.

Our ACE2 mutational scanning showed considerable variability between the three experimental replicates. A potential source of this variability could be the enzyme reaction incubation. We incubate the enzyme reaction for approximately 90 minutes using a ∼30 cm piece of tubing that connects the drop maker chip and the sorting chip. Droplets are generated on the drop maker chip; these droplets then traverse the delay line for ∼90 minutes allowing the enzyme reaction to occur, and are then reinjected onto the sorter chip that reads fluorescence and sorts individual droplets. The laminar flow within the delay line has parabolic velocity profile, creating variation in the incubation time for each droplet. Varied enzyme reaction times will lead to challenges distinguishing ACE2 variants based on their activity. This issue can be addressed in future designs by removing excess oil after drop making to pack the drops together that migrate through the tubing as a plug (controlling droplet incubation using close-packed plug flow).

We found at least five amino acid substitutions at position K187 improve enzyme activity, indicating lysine at this position is suboptimal. K187 is located near the substrate binding site and forms a direct salt bridge with D509, which is adjacent to the Y510 that that forms the S1 active site subpocket. We have previously found mutations at Y510 lead to improvements in ACE2 activity and specificity [6]. Given this observation and the proximity of D509 to Y510, it is reasonable to posit that mutations at position 187 can lead to increased *k*_*cat*_ and decreased *K*_*M*_ by altering the positioning of D509 in ways that change the charge environment around and/or positioning of Y510 within the ACE2 active site [15].

## Conclusions

Our approach discovered ACE2 variants with improved *k*_*cat*_, *K*_*M*_, and *k*_*cat*_/*K*_*M*_. The most important property for therapeutic ACE2 is however, enzyme activity at physiological Ang-II concentrations in the sub-micromolar range. These concentrations are well below the wildtype ACE2’s *K*_*M*_. Our K187 mutants showed improved activity at Ang-II concentrations below 1 μM, indicating potential improvements as a therapeutic. Future ACE2 engineering work should focus on improving catalytic rates at physiological Ang-II concentrations. Although this may be challenging with the current amino acid detection systems due to limited assay sensitivity, the results presented here provide a foundation for increasing ACE2 catalytic rate in the presence of physiological concentrations of Ang-II.

In a more general sense that transcends ACE2, peptidases hold immense potential as enzyme therapeutics due to their ability to precisely regulate biologically active peptides. Their applications span a wide range of therapeutic areas, including the treatment of metabolic disorders [16], cancer [17], infectious diseases [3], and cardiovascular conditions [18] with peptidase therapeutic efficacy being dependent upon stability, activity, and specificity, all of which can vary significantly among different enzyme variants. Protein engineering techniques are essential to tailor these properties to optimize peptidases to meet the stringent requirements of clinical applications, thereby unlocking their full potential as powerful and precise therapeutic agents. Our peptidase library screening method is a valuable addition to the collection of procedures that can be used in the context of such therapeutic development efforts.

