## Additional file 1 for "Droplet microfluidic screening to engineer angiotensin-converting enzyme 2 (ACE2) catalytic activity"

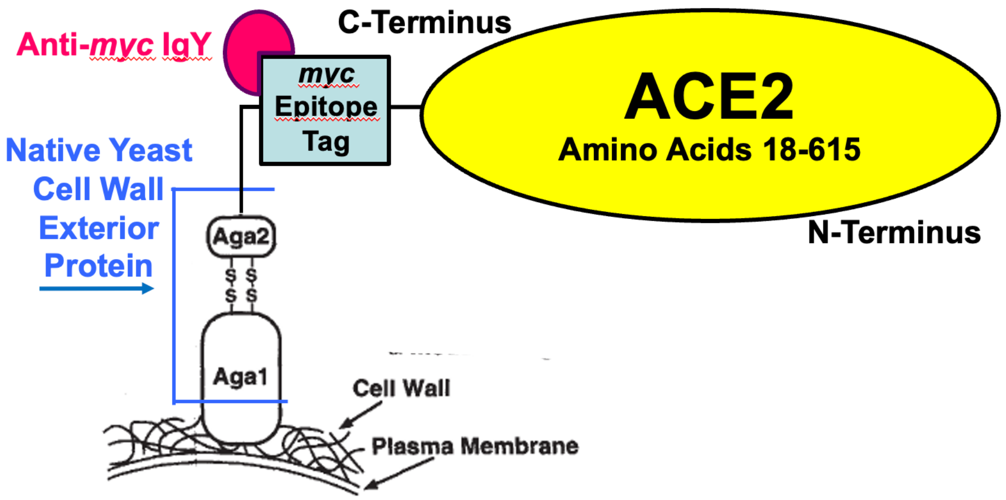


**Figure S1.** Yeast display schematic. ACE2 C-terminus linked to Aga2 native yeast surface protein on yeast cell wall. Anti-*myc* IgY (from chicken) used to quantify ACE2 display. Anti-*myc* IgY is not fluorescently labeled; fluorescent detection of *myc* epitope tag achieved via incubation with Alexa488-conjugated anti-chicken IgG (from goat) as secondary label (not depicted). Each yeast cell displays up to $10^4$ copies of a single ACE2 variant on its surface. Schematic adapted from Boder and Wittrup. *Nat Biotechnol.* 1997 15(6):553-7.

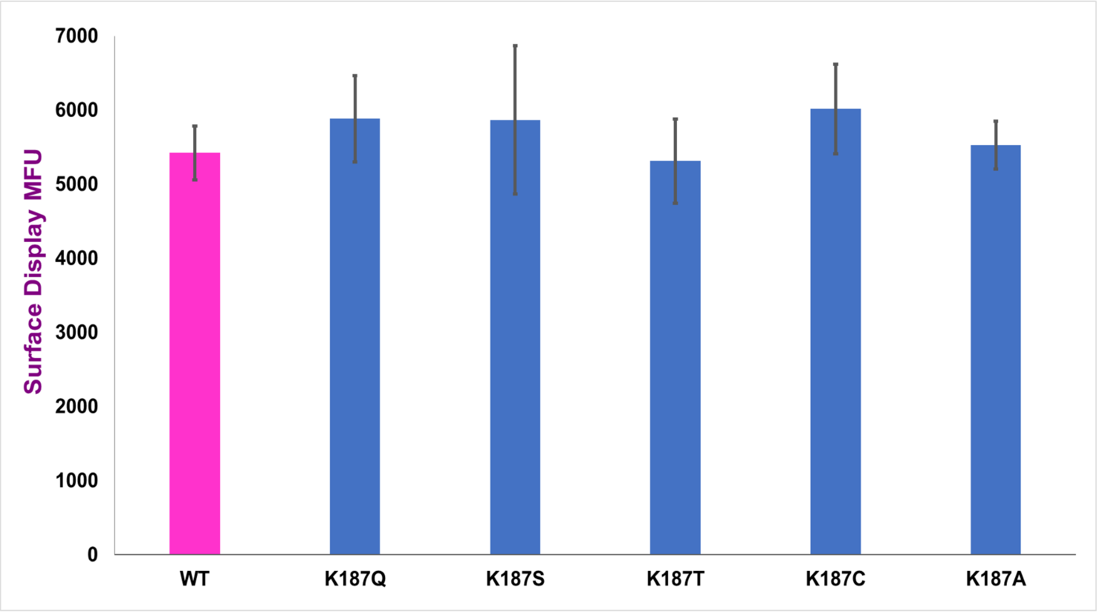


**Figure S2**. Flow cytometry assay mean fluorescence unit values for yeast surface-displayed ACE2s. Error bars in plots denote standard deviations for triplicate measurements.

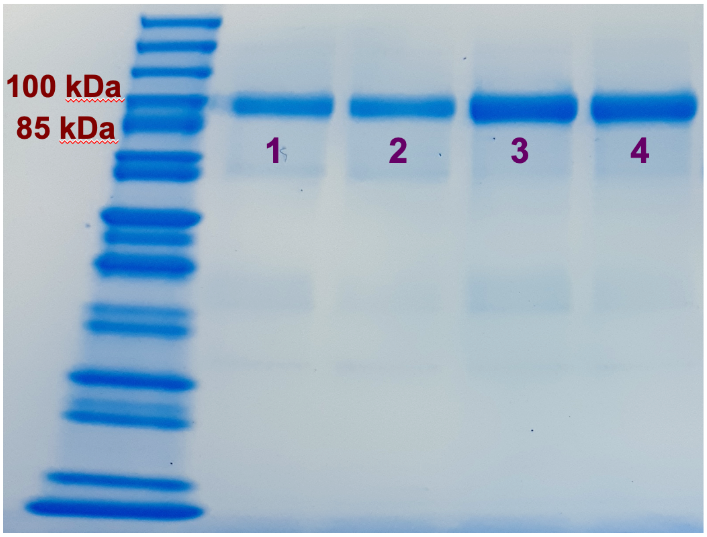


**Figure S3.** Post-purification SDS-PAGE analysis of wild type and K187 mutant ACE2s. Gel lanes (1-to-4) - wt ACE2, K187Q, K187T, and previously characterized [5] T371L/Y510Ile ACE2 mutant that was included as additional comparative standard for K187 mutant ACE2 expression level.
